# Genealogical asymmetry under the IM model and a two-taxon test for gene flow

**DOI:** 10.1101/2024.05.24.595831

**Authors:** Alexander Mackintosh, Derek Setter

## Abstract

Methods for detecting gene flow between populations often rely on asymmetry in the average length of particular genealogical branches, with the ABBA-BABA test being a well known example. Currently, asymmetry-based methods cannot be applied to a pair of populations and such analyses are instead performed using model-based methods. Here we investigate genealogical asymmetry under a two-population isolation-migration model. We focus on genealogies where the first coalescence event is between lineages sampled from different populations, as the external branches of these genealogies have equal expected length as long as there is no post-divergence gene flow. We show that unidirectional gene flow breaks this symmetry and results in the recipient population having longer external branches. We derive expectations for the probability of this genealogical asymmetry under the isolation-migration model and propose a simple statistic (*A_m_*) to detect it from genome sequence data. *A_m_* provides a two-taxon test for gene flow that only requires a single unphased diploid genome from each population, with no outgroup information. We use analytic expectations and coalescent simulations to explore how recombination, unequal effective population sizes and bidirectional gene flow influence *A_m_* and find that the statistic provides unambiguous evidence for gene flow under a continent-island history. We estimate *A_m_* for genome sequence data from *Heliconius* butterflies and *Odocoileus* deer, generating results consistent with previous model-based analyses. Our work highlights a signal of gene flow overlooked to date and provides a method that complements existing approaches for investigating the demographic history of recently diverged populations.

## Introduction

Hybridisation between members of closely related species can sometimes be observed in nature. Depending on the reduction in fitness suffered by early generation hybrids, this process can lead to gene flow between species. Early investigations of the sequence variation within whole genome sequence (WGS) datasets showed that historic gene flow is common, even between species separated by millions of generations of divergence (Kulathinal *et al*. 2009; Martin *et al*. 2013). Over the last decade there has been a considerable increase in the volume of WGS data being generated from natural populations, along with a similar increase in methods for detecting and characterising gene flow (Hibbins and Hahn 2022). As a result, it is now possible to obtain detailed information about past gene flow and ongoing hybridisation within natural systems (see Jensen *et al*. 2023; Marcionetti *et al*. 2024; Satokangas *et al*. 2023 for some recent examples). At the same time, understanding how gene flow shapes patterns of sequence variation, and how this can in turn be used to infer the evolutionary history of populations, is still an active area of research (Galtier 2024; Cousins *et al*. 2024).

### Asymmetry-based methods for detecting gene flow

A natural way to think about how past gene flow can be inferred from WGS data is to consider the genealogical history of individuals sampled from multiple populations. Such a history can be described by the multi-species coalescent (Kingman 1982; Tajima 1983; Rannala and Yang 2003; Jiao *et al*. 2021). Even under a history of strict bifurcation, the stochasticity of the coalescent process results in individual genealogies that do not match the species history (Tajima 1983; Takahata 1989). Such genealogies are generated when multiple lineages with different sampling locations reach the same ancestral population, with the random order of coalescence events generating alternative topologies with equal probability. Post-divergence gene flow, however, leads to certain topologies being more common than others, and, consequently, to an asymmetry in the average length of particular genealogical branches. The ABBA-BABA test is a well known approach for leveraging such genealogical asymmetry to infer gene flow. This four-taxon test was first used to detect historic gene flow between Neanderthals and the ancestors of non-African humans (Green *et al*. 2010), and has since been applied to hundreds of datasets. The test detects past gene flow between non-sister populations, but gives little to no information about the rate, direction, and timing of that gene flow. This has motivated the development of other asymmetry-based methods, that either have improved precision (Martin *et al*. 2015; Lopez Fang *et al*. 2024) or provide more detailed information than the original ABBA-BABA test (Pease and Hahn 2015; Hamlin *et al*. 2020; Martin and Amos 2021). Two major strengths of asymmetry-based approaches are their flexibility and simplicity. For example, the history of gene flow across a sample of tens or even hundreds of populations can be investigated by applying the ABBA-BABA test to all relevant quartets and partitioning the signal across branches of the species tree (Eaton and Ree 2013; Jensen *et al*. 2023). It is also possible to apply asymmetry-based methods across genomes to identify regions with a history of introgression (Malinsky *et al*. 2015; Martin *et al*. 2015). All the while, it is clear what information in the data is used to detect gene flow - asymmetry in polymorphism patterns that are expected to be symmetrical under a coalescent process without gene flow.

Despite the strengths mentioned above, the ABBA-BABA test and other asymmetry-based methods are fundamentally limited by the fact that they only consider a small fraction of the information contained within a sample of genomes (Lohse and Frantz 2014; Martin and Amos 2021). Consequently, these methods cannot distinguish between gene flow and certain scenarios of ancestral population structure (Durand *et al*. 2011) or differences in mutation rate between populations (Xiong *et al*. 2022; Frankel and Ané 2023 but see Koppetsch *et al*. 2023).

### Model-based methods for inferring gene flow

An alternative class of methods are those that explicitly model gene flow within a demographic history (Beerli and Felsenstein 1999; Hey and Nielsen 2004). Given a sample of genome sequences from two or more populations, such methods can be used to calculate statistical support for models with and without gene flow. These methods typically also generate estimates of effective migration rate (*m_e_*), or admixture proportions, along with the other demographic parameters in the model. One attraction of model-based inference is that researchers can obtain a ‘best fitting’ evolutionary history that is (ostensibly) straightforward to interpret, rather than only testing for the presence or absence of gene flow. Additionally, these methods will tend to leverage more of the information contained in genome sequence data than asymmetry-based methods that focus on just a small subset of polymorphism patterns. For these reasons, model-based inference of divergence with gene flow has become common within population genomics, with a wide selection of methods to choose from (e.g. Gutenkunst *et al*. 2009; Rogers 2019; Flouri *et al*. 2020; Excoffier *et al*. 2021). A model-based approach is particularly favourable when the number of populations is small, as sequence variation is straightforward to summarise (e.g. via the joint site frequency spectrum) and the number of alternative models to consider is not too large (although see Dilber and Terhorst 2024).

Given that model-based methods use rich summaries of sequence variation to infer detailed population histories, it is not immediately clear why any researcher would favour an asymmetry-based approach for detecting gene flow, especially if the number of populations is small. However, model-based methods do have several drawbacks. For instance, it is not always straightforward to understand what information in the data provides evidence for gene flow, and more importantly, whether that evidence could instead be the result of bioinformatic artefacts (Shafer *et al*. 2017). The results of model-based inference are also contingent on the finite set of necessarily simple models for which likelihoods can be calculated. If the true demographic history contains processes that cannot be well approximated by simple models, then all models may fit the data poorly (Momigliano *et al*. 2021 but see Huang *et al*. 2022).

Recently, Smith and Hahn (2023) used forward simulations to show that commonly used model-based inference methods can erroneously infer gene flow between fully isolated sister populations when the true history includes natural selection. Although the inferred rates of gene flow were very small, their results highlight the problem of fitting simple demographic models that assume selective neutrality when the evolution of real populations is complex and involves multiple selective forces. Smith and Hahn (2023) suggest that asymmetry-based methods are less likely to lead to false inference of gene flow under pervasive natural selection, mainly because selection should equally affect the length of all genealogical branches in a given population. They point out, however, that asymmetry-based methods cannot currently be applied to datasets containing samples from only two populations.

### Overview

In this work, we investigate how asymmetry in genealogical branch lengths can be used to detect past gene flow between two recently diverged populations. We use the isolation-migration (IM) model as a framework to investigate the properties of genealogies using a small sample of genomes. Our aim, however, is not to use asymmetry in branch lengths to infer parameters of the IM model. We instead aim to highlight genealogical asymmetry as an identifiable signal generated by gene flow under a two-population coalescent process, and we expect our results to be applicable to most other models of population divergence (e.g. secondary contact, migration-only, a single pulse of admixture, etc.).

First, we derive expectations for the length of external branches on genealogies in which the first coalescence event occurs between lineages sampled from different populations. We show that the population-specific external branch lengths of such genealogies are equal in the absence of post-divergence gene flow, but that unidirectional gene flow breaks this symmetry. Next, we present a straightforward approach for estimating this effect from genome sequence data and modify our original derivation to be conditional on the presence of particular mutations. We then use simulations to confirm the accuracy of our analytic results as well as to explore the effects of recombination, unequal population sizes and bi-directional gene flow. Finally, we use our estimator of genealogical asymmetry as a two-taxon test for gene flow and apply it to two real-world examples with contrasting demographic histories.

## Results

### Genealogical incongruence and asymmetry

We focus on an IM model with five parameters (Figure 1A). The parameters *N_A_*, *N_B_* and *N_AB_* correspond to the effective sizes of populations *A*, *B* and the ancestral population, respectively. The parameter *t* is the number of generations between the present and the onset of divergence, and *m_e_*is the effective rate of migration (pastward) between populations *A* and *B*. Initially, we only consider unidirectional migration, with *m_e_*representing the rate at which lineages migrate from population *A* to population *B* backwards in time, with no migration in the opposite direction.

**Figure 1:**
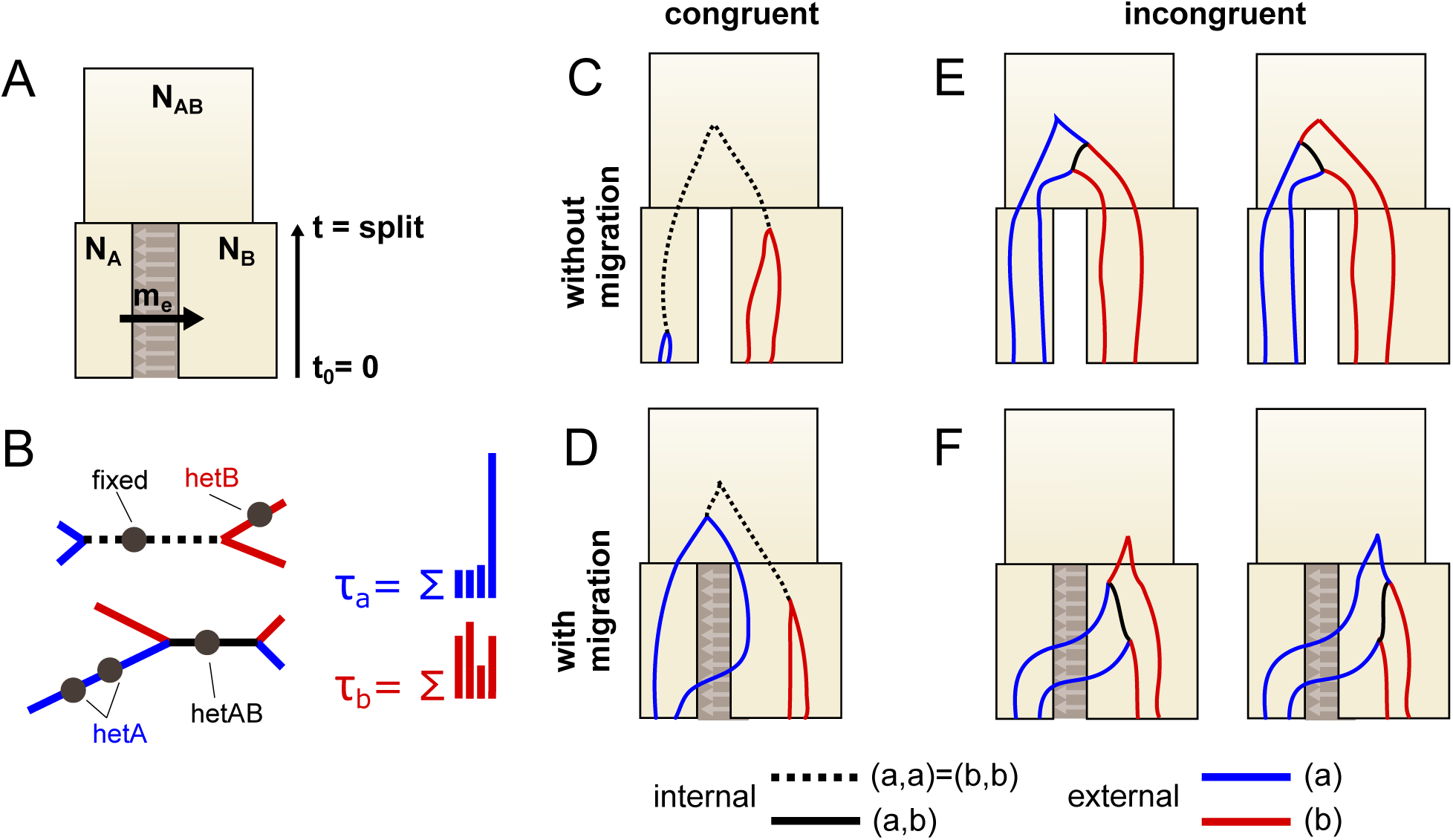
(**A**) The five-parameter IM model includes the sizes of the sampled populations *N_A_* and *N_B_*, an effective (backward) migration rate *m_e_*, a population split time *t*, and the ancesteral population sizes *N_AB_*. (**B**) Unrooted genealogies may be congruent, having an internal (a,a)/(b,b) type branch (dashed black), or incongruent, having an internal (a,b) type branch (solid black). External branches (a) in blue and (b) in red distinguish samples from the *A* and *B* populations respectively. *τ_A_* and *τ_B_*are the total branch length of (a) and (b) type branches, respectively. In polymorphism data, only mutations on the internal branches can identify the topologies as congruent (*fixed*) or incongruent (*hetAB*), and the relative length of external branches is reflected in the number of *hetA* and *hetB* type mutations. (**C** and **D**) Congruent genealogies are generated whenever the first coalescence event is between lineages sampled from the same population. (**E** and **F**) Incongruent genealogies, where the first coalescence event is between lineages sampled from different population, can be generated through failed coalescence (**E**) or migration (**F**).

We consider a sample of two lineages (equivalently, a diploid) from each population (Figure 1). Genealogies from this sampling scheme contain four leaves, and if leaves are labelled by the population from which they are sampled (either *a* or *b*), there are two possible unrooted topologies: those in which the first coalescence event is *a* + *a* or *b* + *b* (congruent to the species tree), and those in which the first coalescence event is *a* + *b* (incongruent) (Figure 1B). We identify branches on these genealogies as either external or internal, with external branches defined as those leading directly to a leaf on the unrooted genealogy. Labelling branches by their descendants results in external branches with the labels *a* and *b*, and internal branches with the labels *aa* (equivalent to *bb* on an unrooted tree) and *ab* (Figure 1B). The total length of branch types (*τ_a_*, *τ_b_*, *τ_aa_*, *τ_ab_*) are (non-independent) random variables with a complicated joint distribution determined by the coalescent process. For simplicity, we focus only on the marginal distributions of the total external branch lengths (*τ_a_* and *τ_b_*), particularly their expected values 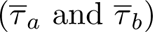.

Under the IM model, the total (marginal) length of external branches *a* and *b* (*τ_a_* and *τ_b_*) depend on all five parameters, and in general, we do not expect 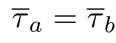. Equality of the expected branch lengths only occurs under two scenarios: when there is no divergence (i.e. *t* = 0) or when there is divergence without gene flow between two equal-sized populations (i.e. *t >* 0 and *m_e_* = 0 with *N_A_* = *N_B_*). In other words, for any biologically realistic case of population divergence we expect 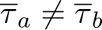 (Figure 1C and D). However, if one focuses exclusively on genealogies with an incongruent topology (Figure 1E and F), 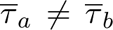 is an unambiguous indicator of post-divergence gene flow between the two populations. We can understand this as follows.

Incongruence can be generated two ways: either failed coalescence in populations *A* and *B* or through gene flow. Assuming no gene flow (*m_e_* = 0), incongruence occurs solely due to failed coalescence and with probability

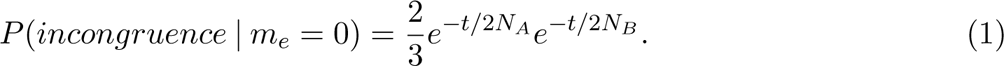

For such genealogies, all four lineages must be present in the ancestral population and the order in which they coalesce is random (Figure 1E). Conditioned on incongruence, the expected value of the (marginal) total branch length distributions is

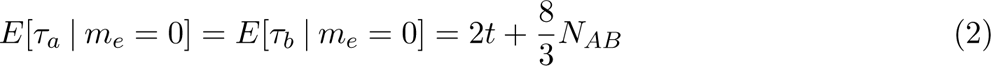

The fact that Equation (2) applies to both *a* and *b* branches and is independent of *N_A_* and *N_B_*means that, on-average, genealogies with an incongruent topology are expected to have equal external branch lengths 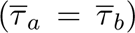 when *m_e_*= 0 even if populations *A* and *B* have drastically different effective population sizes (Figure 1E). Although this result is not particularly surprising, it is important in that it provides an expectation for symmetry in external branch lengths (conditional on incongruence) under a two-population history without gene flow.

Given the result above, we next investigate whether post-divergence gene flow breaks or maintains symmetry in external branch lengths. For simplicity, we now assume that all three populations share the same *N_e_* and use the parameters *M* = 2*N_e_m_e_* and *T* = *t/*2*N_e_* for ease of notation. Lohse *et al*. (2011, 2016) derived the distribution of branch lengths for genealogies with small sample size under the IM model. Here we use the same approach – the generating function (GF) of genealogical branches (Lohse *et al*. 2011) – but focus on incongruent genealogies. The GF is a series of terms, each representing a unique sequence of coalescence, migration, and population split events. We refer to these as the *paths* of the IM model, of which there are 95. For example, the path for the genealogy in Figure 1D is defined by the following events (looking pastward in time): migration from *A* to *B*, coalescence of the *b* lineages, the population split, coalescence of the *a* lineages, and finally coalescence of the *aa* and *bb* lineages. By extracting the 33 terms in the GF that correspond to paths with an incongruent topology (i.e. those that contain *ab* branches), the probability of incongruence can be calculated as:

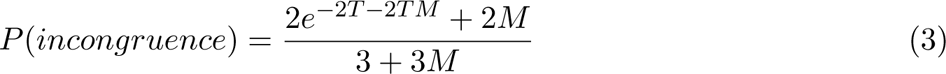

Setting *M* = 0 recovers Equation (1), while *M >* 0 yields a higher probability of observing an incongruent genealogy.

We can gain information about how external branch lengths are affected by gene flow by obtaining the expected values of *τ_a_* and *τ_b_* independently for each of the 33 incongruent paths in the GF. Of these paths, 10 have expected branch lengths with 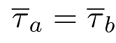. Another 20 have expected branch lengths where 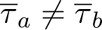, but each is an equiprobable member of a pair, for which the combined branch lengths of *a* and *b* are equal (e.g. Figure 1E and F). This is because the members of each pair are exchangeable but for the order of the last two coalescence events. The three remaining paths are unpaired and have expected branch lengths 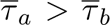. It is these three paths, shown in Figure 2A, that break the symmetry in external branch lengths on incongruent genealogies when *M >* 0. We hereafter refer to these genealogies as asymmetrical. In the same figure, we show the probability of each such genealogical history (Figure 2B) as well as the expected values of *τ_a_* and *τ_b_* (Figure 2C) in relation to *M* (here, assuming *T* = 1). These results show that the probability of asymmetrical genealogies peaks at intermediate values of *M* (Figure 2B), but that the difference in 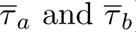 remains substantial across *M* values (Figure 2C). Note as well that these asymmetrical incongruent genealogies are only observed when there is gene flow, i.e. they have zero probability when *M* = 0 (Figure 2B).

**Figure 2:**
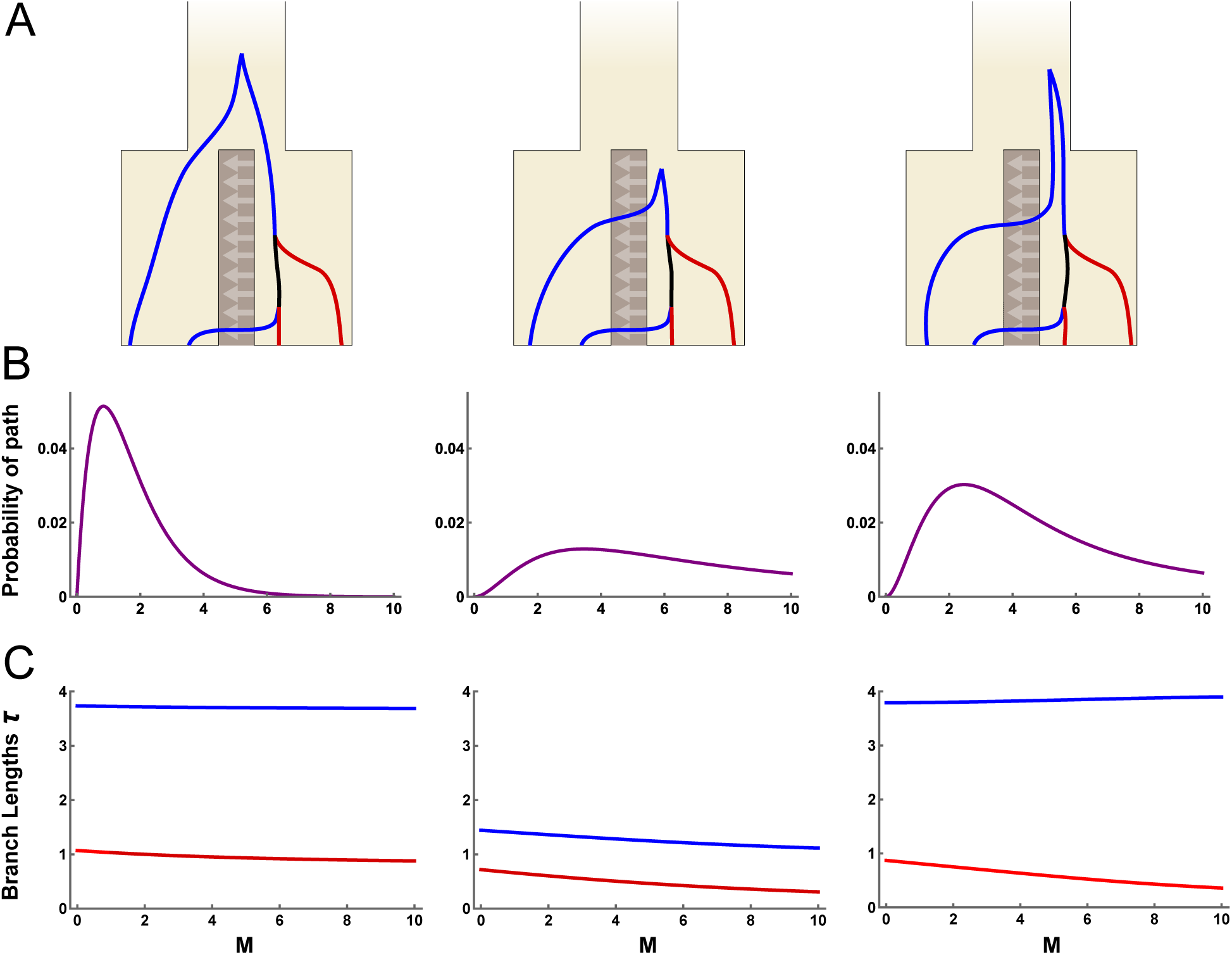
The three asymmetrical incongruent genealogies (**A**) are generated when the first event is migration, the second event is (a)+(b) coalescence, and the third event is (a, b)+(b) coalescence with the remaining (a) lineage still isolated in population *A*. Here, blue and red correspond to the external branches for samples from the *A* and *B* population, respectively (see Figure 1). Each genealogy has a unique probability of occurring (**B**) and a different disparity in the expected total length of external branches (**C**), both of which depend on the split time (here, *T* = 1.0) and the migration rate *M* (x-axis). Branch lengths *τ_a_* and *τ_b_* are shown in blue and red, respectively, and in units of 2*N_e_* generations. In all plots *N_A_* = *N_B_* = *N_AB_*.

The above results suggest that, across incongruent genealogies, the asymmetry of *a* and *b* branch lengths can be used to distinguish between divergence with and without gene flow. Additionally, the relative values of *τ_a_* and *τ_b_* also provide information about the direction of gene flow, as the recipient population (forwards in time) is expected to have longer external branches. We next calculate values of *τ_a_*and *τ_b_*conditional on sampling a random incongruent genealogy, rather than a specific path. Figure 3 shows the probability of incongruence as well as the 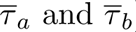 of incongruent genealogies, dependent on *M* and *T* . Additionally, we summarise the difference in external branch lengths using the scaled ratio

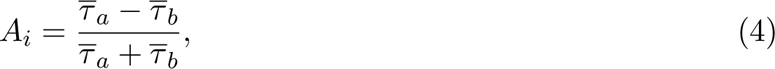

**Figure 3:**
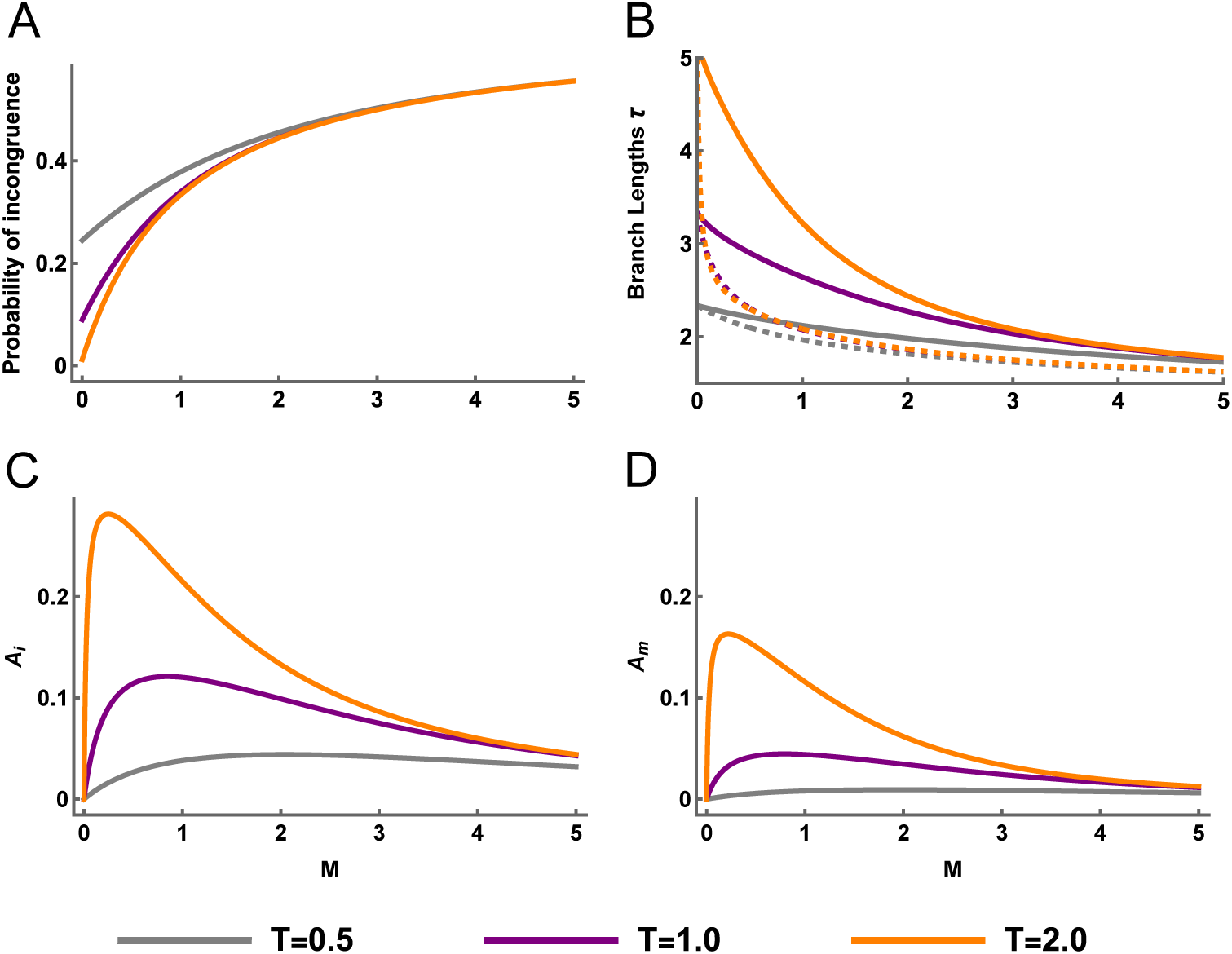
The effect of migration rate *M* and divergence time *T* on (**A**) the probability of incongruence, (**B**) the expected total length of external branches, (**C**) the scaled difference in external branch lengths (*A_i_*) and (**D**) the scaled difference in external branch lengths conditional on observing a *hetAB* mutation in a 200 bp block of sequence (*A_m_*), given a mutation rate of 1 *×* 10*^−^*^8^ per-site per-generation and an *N_e_* of 100, 000 in all populations. Grey, purple, and orange lines correspond to divergence times *T* = *{*0.5, 1.0, and 2.0*}*, respectively. In panel **B**, the solid lines correspond to *τ_a_*, the dashed lines to *τ_b_*, and branch lengths are measured in units of 2*N_e_*generations.

where 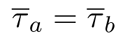 leads to 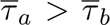 to *A_i_ >* 0 and 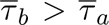 to *A_i_ <* 0. Intermediate values of *M* generate the greatest inequality in *τ_a_* and *τ_b_*, and therefore large values of *A_i_*(Figure 3C). This can be explained by the fact that asymmetrical genealogies are only generated when just one lineage migrates before the first coalescence event (Figure 2). As the divergence time *T* increases, so does *A_i_* (Figure 3C). With no divergence (*T* = 0) all four lineages are effectively sampled from the ancestral population and so *A_i_* = 0. By contrast, a migration only model (*T* = *∞*) eliminates the possibility that the first event is the merging of populations *A* and *B*, thereby maximising both the frequency of asymmetrical genealogies and *A_i_*.

For the biologically relevant range of parameters we consider in Figure 3, we observe a maximum *A_i_* of 0.282. In principle, *A_i_* can be as high as 1.0, but such high values are only expected when *T* is large and *M* is small. For example, when *T* = 10 and *M* = 1 *×* 10*^−^*^4^, *A_i_* = 0.775, but the probability that any genealogy is incongruent is extremely low (6.7*×*10*^−^*^5^). With so few incongruent genealogies from which to estimate *A_i_*, we do not expect to be able to use *A_i_* for inference under this demography. However, we expect *A_i_* to be a useful summary statistic for detecting gene flow between recently diverged populations where incongruent genealogies are common (Figure 3).

### Estimating asymmetry from patterns of mutation

Given that asymmetry in external branch lengths on incongruent genealogies provides evidence for historic gene flow, how can we identify such genealogies in genome sequence data? Put differently, how can we estimate *A_i_*? One approach would be to reconstruct the ancestral recombination graph (ARG) for a sample of genomes, and then summarise external branches across the ARG to calculate *A_i_*. While possible (see Discussion), we instead investigate a methodologically simpler approach that uses only a single diploid sample from each population and requires neither phase information nor polarization.

For this sampling scheme (an unphased and unpolarised diploid from each population), there are four types of polymorphisms corresponding to the distinct branches of the genealogy: *hetA*, *hetB*, *hetAB*, *fixed* (here, following the nomenclature of Laetsch *et al*. 2023). Assuming an infinite sites mutation model, a *hetAB* polymorphism (in which both diploids are heterozygous for the same alleles) can only be generated when the underlying genealogy is incongruent (Figure 1B). These mutations mark the approximate locations of a subset of the incongruent genealogies throughout the genome, and we can modify our estimator of genealogical asymmetry by restricting ourselves to those loci. We define

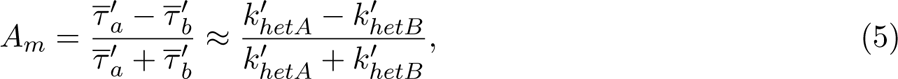

where 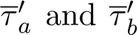 are the expected external branch lengths of genealogies containing at least one mutation on the *ab* branch. This can be approximated by the counts *k^′^* of *hetA* and *hetB* polymor-phisms that lie within *l* bases of a *hetAB* polymorphism, where 2*l* is chosen to be small enough to limit the probability that the region spans more than one genealogy.

In the previous section, we defined *A_i_* as a ratio between the expected *a* and *b* branch lengths of incongruent genealogies. *A_m_*instead estimates the ratio of *a* and *b* branch lengths in the context of genealogies with at least one *hetAB* polymorphism. This is a non-random subset of incongruent genealogies because those with longer internal branches are more likely to carry a *hetAB* mutation. We therefore expect values of *A_i_*and *A_m_*to be highly correlated but to differ in scale (Figure 3D). To resolve this discordance, we again use the GF approach and derive the expected branch lengths 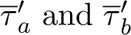 (and therefore *A_m_*), this time conditioning on those genealogies with at least one *hetAB* mutation. *A_m_*now depends on the rate of mutation on the *ab* branch which is parameterised by a per-site mutation rate *µ* and locus length 2*l*. The GF conditioned on observing *≥* 1 *hetAB* mutations is obtained by (i) deriving the GF over which the number of *hetAB* mutations is marginalized (that is, accounts for any number of *hetAB* mutations) then (ii) subtracting the GF conditioned on observing exactly *k_hetAB_* = 0 mutations. For details see the supplementary Mathematica notebook. We used coalescent simulations to confirm the accuracy of our derivations of *A_i_* and *A_m_*. Specifically, we simulated divergence with gene flow under IM models with parameters *N_A_* = *N_B_* = *N_AB_* = 100, 000, *T* = 1.0, *M ∈ {*0, 1.0, 3.0*}* and *µ* = 10*^−^*^8^. We simulated 500, 000 non-recombining sequences of size 2*l* = 200 and sampled a single diploid from each population. The values of *A_i_* and *A_m_* estimated from the branch lengths of simulated genealogies closely match to our analytic expectations (Table 1). Reassuringly, the 95% confidence intervals (95% CIs) for estimates of *A_m_* do not span zero when *M >* 0, showing that the effect of gene flow on genealogical asymmetry can be distinguished from a null-scenario given sufficient information about branch lengths.

**Table 1:** Estimates (est.) of *A_i_* and *A_m_* calculated from simulated genealogies are presented for three different demographic histories. Analytic predictions (pre.) are also given for the same demographic histories. Each estimate is derived from 500,000 simulation replicates of a 200 bp sequence without recombination. The three demographic histories share all parameters (*N_A_* = *N_B_* = *N_AB_* = 100, 000, *T* = 1.0 and *µ* = 10*^−^*^8^) except the rate of migration (*M ∈ {*0, 1.0, 3.0*}*). For simulation derived estimates of *A_i_* and *A_m_*, 95% CIs are given in brackets and were estimated through block jackknife resampling.

| $M$ | est. $A_i$ (95% CIs) | pre. $A_i$ | est. $A_m$ (95% CIs) | pre. $A_m$ |
| --- | --- | --- | --- | --- |
| 0.0 | -0.00046 (-0.00386, 0.00293) | 0.00000 | -0.00190 (-0.00682, 0.00302) | 0.00000 |
| 1.0 | 0.12109 (0.11865, 0.12352) | 0.12016 | 0.04260 (0.03865, 0.04656) | 0.04394 |
| 3.0 | 0.07569 (0.07299, 0.07838) | 0.07494 | 0.02180 (0.01766, 0.02594) | 0.02432 |

### Recombination and unequal effective population sizes

So far, our analytic and simulated results have assumed that a short block of sequence is associated with a single genealogy. In reality, short blocks of sequence centered on a *hetAB* polymorphism will sometimes reflect multiple genealogies due to recombination, and this has the potential to bias our summary statistic (*A_m_*) away from its expected value. This is because recombination violates the assumption that nearby *hetA* and *hetB* polymorphisms are on the same incongruent genealogy as the focal *hetAB* polymorphism. Estimates of *A_m_* in the limit of high recombination would reflect the average asymmetry in external branch lengths across all genealogies rather than just those that are incongruent, and it would no longer be a valid test for gene flow. It is less clear, however, what to expect when calculating *A_m_* from short blocks of sequence (i.e. a few hundred bases) in which recombination only happens occasionally.

To understand the effect of recombination on estimates of *A_m_* we simulated population divergence with gene flow as in Table 1 with *M* = 1.0 but with the addition of recombination at a rate of *r* = 10*^−^*^8^ per-base per-generation. We simulated sequences of 66 kb (5000 replicates) and estimated *A_m_* from genealogical branch lengths using different block sizes and therefore varying frequencies of recombination. We find that simulated values of *A_m_* closely match the analytic expectation (which assumes no recombination) for very small blocks but then diverge to greater values once the block size reaches around 100 bases (Figure 4A). At large block lengths, e.g. 16 kb, estimates of *A_m_* reach the expectation for asymmetry in external branch lengths given a random sampling of genealogies. These results show that too large a block length can lead to the inclusion of congruent genealogies through recombination and therefore biased estimates of *A_m_*.

**Figure 4:**
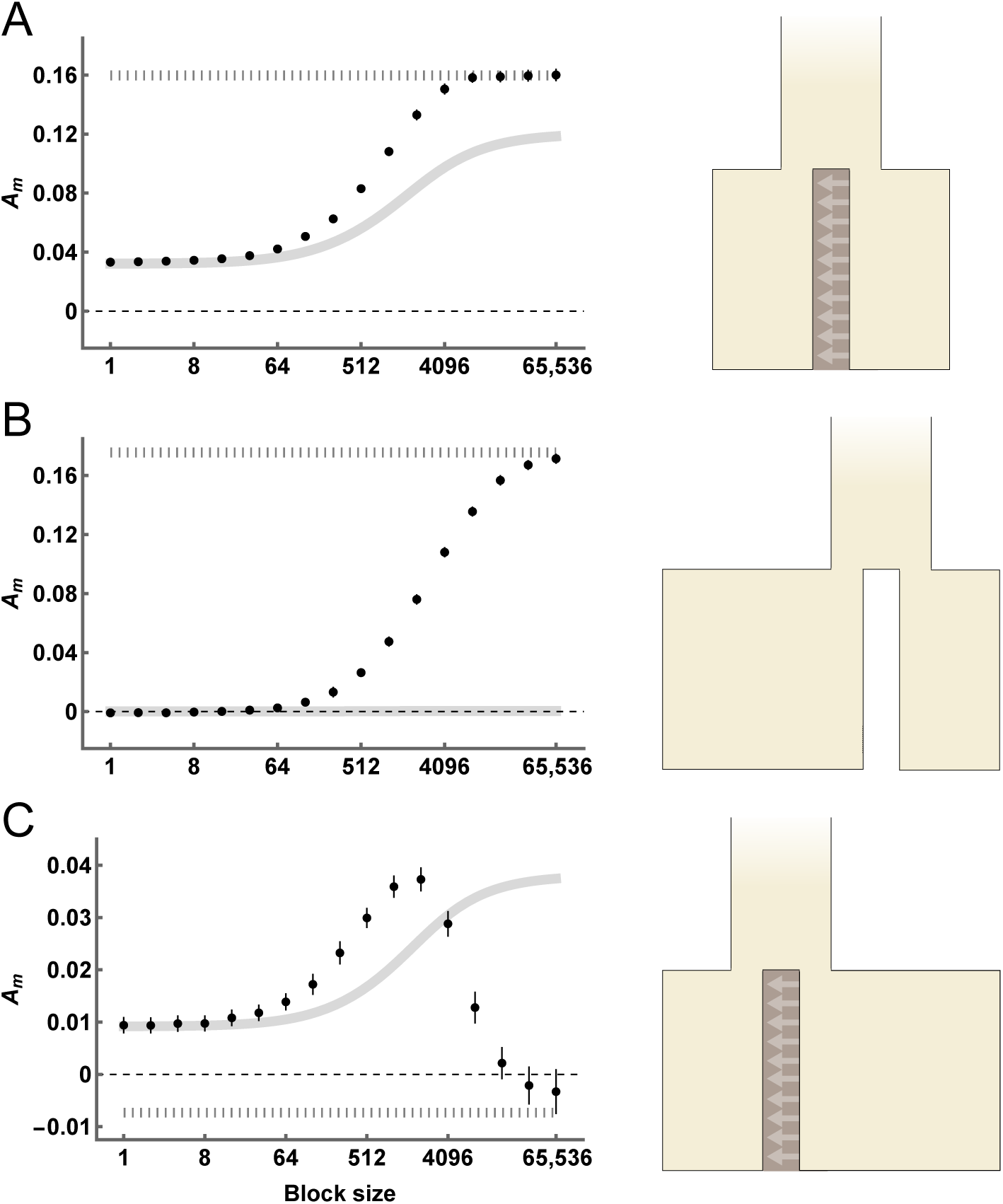
Estimates of *A_m_* for various block sizes for simulations with recombination and a demographic model with (**A**) Equal population sizes *N_A_* = *N_B_* = *N_AB_* = 100, 000 and unidirectional migration at rate *M* = 1.0 from population *B* to *A* (forwards in time), (**B**) Unequal population sizes *N_A_* = 200, 000 and *N_B_* = *N_AB_* = 100, 000 with strict isolation *M* = 0, and (**C**) Unequal population sizes *N_A_* = *N_AB_* = 100, 000 and *N_B_* = 200, 000 with unidirectional migration from *B* to *A* forwards in time at rate *M* = 1.0. In all panels the divergence time *T* = 1.0, the per-base mutation rate *µ* = 10*^−^*^8^, and the recombination rate *r* = 10*^−^*^8^ per-base per-generation. The analytic expectation for *A_m_* in the absence of recombination is shown in a thick solid line, whereas the analytic expectation for *A_m_* under free recombination is shown as a thick dashed line. Error bars correspond to the 95% CIs of each estimate.

Next, we simulated a history of divergence with strict isolation (*M* = 0), but with inequality in *N_e_* between the sampled populations (*N_A_*= 200, 000 and *N_B_* = 100, 000). We performed the same analysis as above and find that *A_m_* = 0 at short block sizes, as expected under a history of strict isolation (Figure 4B). However, as the block size increases *A_m_* becomes significantly greater than zero and eventually reaches the value expected for random sampling (Figure 4B). In this case, recombination and unequal *N_e_* leads to a false-positive signal of gene flow. This is driven by the inclusion of congruent genealogies, via recombination, where external branch lengths are determined by *N_A_* and *N_B_* (Figure 1C). Although this risk can be minimised by choosing a very short block length, thus reducing the frequency of recombination within blocks, this also reduces the number of linked polymorphisms and therefore the power to detect asymmetry in branch lengths. Recombination in the presence of unequal *N_e_* therefore represents a challenge for detecting gene flow through asymmetry alone.

Finally, we consider a demographic history where *A_m_* is expected to be positive in the absence of recombination, due to gene flow (*M* = 1.0), but *A_m_* in the limit of high recombination is expected to be negative due to population *B* being larger in effective size (*N_A_* = *N_AB_* = 100, 000 and *N_B_*= 200, 000). At very small and very large block sizes, simulations match these expectations (Figure 4C). Surprisingly, the transition between these limits is non-monotonic with an initial increase in *A_m_* after increasing blocks to a few hundred bases in length. We hypothesise that this is because a small number of recombination events reduces the effect of conditioning of a *hetAB* polymorphism (see Figures 3C and 3D), thus increasing *A_m_*, and that this outweighs the introduction of congruent genealogies which decrease *A_m_* in this case.

Overall, these simulations show that recombination can have a significant influence on our measure of incongruent asymmetry. Given that non-zero values of *A_m_* can be generated through recombination despite no post-divergence gene flow, we suggest that *A_m_* always be calculated across a range of block sizes and that the interaction between asymmetry and recombination be interpreted carefully. In the presence of recombination, evidence for post-divergence gene flow can be seen in values of *A_m_* that are stable and consistently non-zero across small block sizes (Figure 4A). The pattern in Figure 4C constitutes even stronger evidence for gene flow, as non-zero values of *A_m_* at small block sizes cannot be explained by the introduction of congruent genealogies via recombination, which would instead decrease *A_m_*. This pattern is an expected consequence of a continent-island demographic history where the population with smaller *N_e_* receives gene flow forwards in time.

### Bidirectional gene flow

In the above sections we have shown how undirectional gene flow generates asymmetry in the external branch lengths of incongruent genealogies. The assumption of strictly unidirectional gene flow allows exact derivations of branch lengths under the IM model using generating functions. It is, however, more challenging to obtain analogous derivations for a model in which gene flow happens in both directions (see Lohse *et al*. 2016). We are nonetheless interested in whether asymmetry is generated under bidirectional gene flow. We therefore use coalescent simulations to estimate expectations for *A_m_* while allowing gene flow in both directions. We focus on the demography in Figure 4A and vary the rate of gene flow in both directions, simulating short sequence blocks of 200 bases without recombination. Our simulations show that even a low rate of opposing gene flow can have a large effect on *A_m_* (Figure 5). At the same time, we find that *A_m_* is typically non-zero under the demographic parameters we considered, and is only indistinguishable from zero when *M_A_ ≈ M_B_*. Additionally, *A_m_* tends to be positive when *M_A_ > M_B_* and negative when *M_B_ > M_A_* (Figure 5), suggesting that *A_m_* provides information about the dominant direction of gene flow, at least in the case of similar effective population sizes. These simulation results also show that the total amount of gene flow (i.e. *M_A_* + *M_B_*) affects the magnitude of *A_m_*. In particular as *M_A_* + *M_B_* becomes large any difference between them is only weakly reflected in *A_m_*. This mirrors the analytic results for unidirectional gene flow (Figure 3D) because both are driven by the fact that very high migration rates lead to a coalescent process similar to that of a panmictic population. Overall, these results show that the power of *A_m_* to detect gene flow is reduced, but not removed, by bidirectional gene flow.

**Figure 5:**
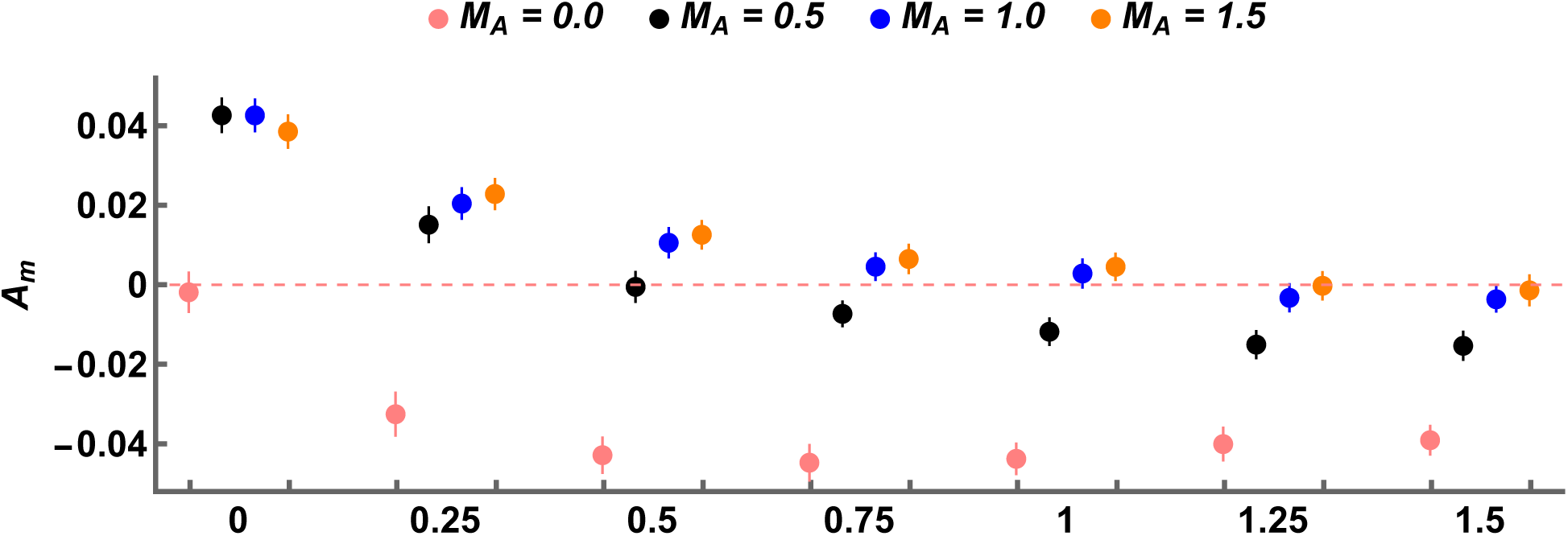
Expected values of *A_m_* obtained by coalescent simulation under an IM model with gene flow in both directions (500,000 replicates per-estimate). All simulations share the parameters *N_A_* = *N_B_* = *N_AB_* = 100, 000, *µ* = 10*^−^*^8^ per-site per-generation, and *T* = 1.0. Simulations vary in the rate of migration from population *A* to population *B* (backwards in time) (*M_A_*, denoted by colour), as well as the rate of migration in the opposite direction (*M_B_*, discrete x-axis). Error bars correspond to the 95% CIs of each estimate.

### Detecting gene flow between *Heliconius* butterflies

To demonstrate how genealogical asymmetry can be used to detect historical gene flow between two populations, we re-analyse genome sequence data from a pair of closely related butterfly species: *Heliconius melpomene* and *H. cydno*. These species exhibit strong assortative mating, and the rare natural hybrids are fertile only when male. Despite this, previous analyses of genome sequence data have revealed evidence for post-divergence gene flow from *H. cydno* to *H. melpomene* forwards in time (Martin *et al*. 2013; Kronforst *et al*. 2013; Lohse *et al*. 2016; Laetsch *et al*. 2023). We calculated *A_m_* across a range of block sizes for a pair of male individuals (one sampled from each species and both from Panama). At short block sizes, *A_m_* is positive (Figure 6A). As the block size increases, however, *A_m_* eventually decreases and reaches negative values (Figure 6A). This matches the pattern observed in Figure 4C and constitutes strong evidence for gene flow from *H. cydno* to *H. melpomene*.

**Figure 6:**
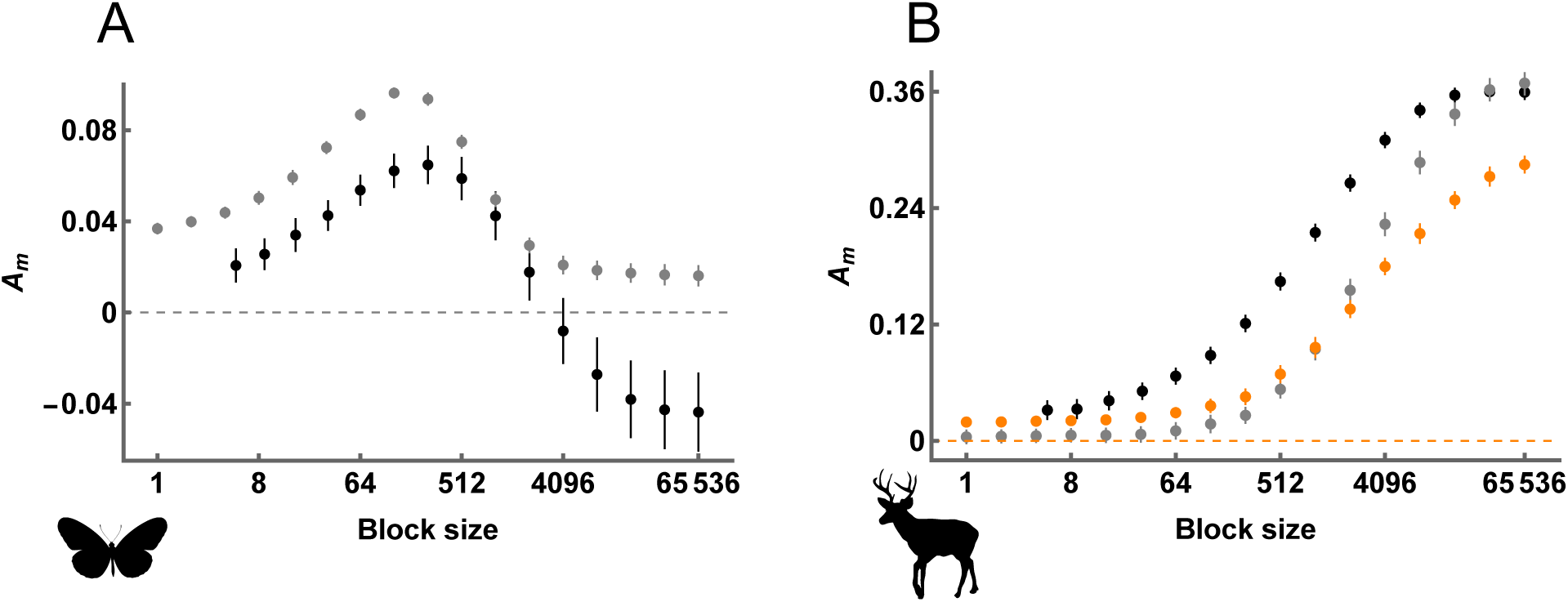
Distributions of *A_m_* (y-axis) across block sizes (x-axis, *log*2 scale) for (**A**) *Heliconius melpomene* and *H. cydno*, and (**B**) *Odocoileus virginianus* and *O. hemionus*. In both panels black points correspond to values of *A_m_* estimated directly from WGS data. Grey points in **A** correspond to values of *A_m_* expected under the demographic history inferred by Laetsch *et al*. (2023). Grey points in **B** correspond to values of *A_m_* expected under the demographic history inferred by Kessler and Shafer (2024) and presented in their Figure 4A. Orange points correspond to values for the alternative history presented in Figure 4B of Kessler and Shafer (2024). Error bars correspond to the 95% CIs of each estimate.

We can also compare our *A_m_*values to those expected under the demographic history of *H. melpomene* and *H. cydno* as inferred by Laetsch *et al*. (2023), whose data we reanalysed. Interestingly, simulating under the inferred demography generates *A_m_* values that only weakly match those from the data (Figure 6A). This is somewhat surprising given that Laetsch *et al*. (2023) fit their model to blockwise data that is informative about the joint distribution of genealogical branch lengths, and therefore contains information about *A_m_* (see Discussion). The discrepancy could be because the approach of Laetsch *et al*. (2023) assumes no recombination within short sequence blocks, as well as an IM demography with unidirectional gene flow. Although our analytic results have the same assumptions, summarising *A_m_* across a range of block sizes does not require any assumptions about recombination or demography.

### Detecting gene flow between *Odocoileus* deer

We expect our approach for detecting gene flow to perform particularly well for the pair of *Heliconius* species analysed above, as the long-term rate of gene flow is intermediate (*M ≈* 1) and the smaller *N_e_* population receives migrants forwards in time (Kronforst *et al*. 2013; Lohse *et al*. 2016). Using the same approach to detect low levels of bidirectional gene flow between sister taxa is likely more challenging. As an example, we re-analyse genome sequence data from white-tailed and mule deer: *Odocoileus virginianus* and *O. hemionus*. Although hybridisation is observed between present-day populations, Kessler *et al*. (2023) showed that the distribution of coalescence times within and between these species (inferred with MSMC-IM; Wang *et al*. 2020) is consistent with a history of speciation without gene flow. The same authors recently expanded their dataset and fit demographic models to the joint site frequency spectrum (jSFS) with dadi (Gutenkunst *et al*. 2009), inferring low levels of gene flow via secondary contact between the two species (Kessler and Shafer 2024). We calculated *A_m_* across a range of block sizes for a pair of individuals with high coverage genome sequences data (Ov ON6 and Oh WA1). We find that *A_m_*is positive when blocks are short and monotonically increases with block length (Figure 6B). Because we cannot estimate *A_m_* in the complete absence of recombination, it is difficult to tell whether these results are evidence of a demographic history with post-divergence gene flow (Figure 4A), or a history without gene flow but a large difference in *N_e_* between the two populations (Figure 4B).

We next perform simulations under two alternative demographic histories inferred by Kessler and Shafer (2024). The first of these demographies includes a low rate of bidirectional secondary contact gene flow, where the probability that a single lineage migrates at least once is low (*P* = 0.015). Expected *A_m_* values for this demography are indistinguishable from zero at small block sizes (grey points in Figure 6B; *A_m_* = 0.0037 (-0.0021, 0.0095) for a block size of 1 base). The second demography includes bidirectional gene flow between the present and the onset of divergence and lineages have a greater probability of migration (*P* = 0.044). In this case, simulated *A_m_* values are greater than zero across all block sizes (orange points in Figure 6B; *A_m_* = 0.0163 (0.0091, 0.0234) for a block size of 1 base). We find that *A_m_* values estimated directly from the data are greater than those of both simulated demographies at small block sizes (Figure 6B). This simulation check therefore suggests that there is indeed some evidence for historic gene flow from *O. hemionus* to *O. virginianus*, forwards in time, even if the signal is subtle overall.

## Discussion

Understanding how past demography shapes patterns of genome sequence variation is a major goal of evolutionary genetics. In this work, we have shown that gene flow between two recently diverged populations leads to an asymmetry in the length of external branches on genealogies that are incongruent with the species history. This asymmetry arises when (i) lineages that were sampled from the same population become trapped in different ancestral populations by a single migration event, (ii) the final coalescent event involves the isolated lineage, and (iii) this coalescence uniquely occurs only after a second migration or a merging of populations (Figure 2A). Importantly, we have shown that this effect is identifiable from WGS data, and we used this to define an asymmetry-based test for gene flow that requires data from only two taxa.

Past gene flow is also expected to influence population genomic analyses beyond tests for gene flow. For example, when inferring population size histories using the distribution of pairwise coalescent times (Li and Durbin 2011), the trapping of lineages in different ancestral populations leads to upwardly biased estimates of *N_e_* (Mazet *et al*. 2015; Cousins *et al*. 2024). This process may also affect inference methods for detecting selective sweeps that are based on local distortions in genealogical branch lengths (Nielsen *et al*. 2005; Bisschop *et al*. 2021). One possibility is that genealogies with long external branches generated by gene flow may resemble a locus at intermediate distance from a beneficial mutation and generate false sweep signatures . It has also recently been shown that gene flow can bias estimates of the recombination rate from inference methods based on measures of linkage disequilibrium (Setter *et al*. 2022; Samuk and Noor 2022). Although not our focus, estimation of *A_m_* may then be a useful way to check whether a given dataset meets the assumption of panmixia, which many population genomic inference methods require.

### Comparison with other approaches

Our approach to detecting gene flow shares several similarities with the four-taxon ABBA-BABA test (Green *et al*. 2010; Durand *et al*. 2011). Both focus on incongruent genealogies, and both rely on the expectation that particular branches are (on average) equal in length under a history of strict isolation but unequal under a history of gene flow. They also both share the general strengths of asymmetry-based methods: they require few assumptions about the demographic history and are expected to be robust to the effects of natural selection. There are, however, several important differences between the two approaches. For one, *A_m_* provides information about the direction of gene flow (Figure 5), whereas the ABBA-BABA test does not (although see Pease and Hahn 2015 and Leppäaläa *et al*. 2024 for five-taxon extensions that do). Additionally, the ABBA-BABA test statistic (Patterson’s D) typically scales with the rate of gene flow, whereas *A_m_*has a non-linear and non-monotone relationship with *M* (Figure 3). Another key difference is that the ABBA-BABA test assumes independence among all sites and uses information about the average length of genealogical branches. By contrast, *A_m_* leverages linkage information to estimate the average lengths of branches for genealogies that carry at least one mutation on an incongruent internal branch. As we do not know the true span of any genealogy, we can only take these measurements in small regions that we (optimistically) assume are non-recombining. So, while *A_m_*shares many of the strengths of other asymmetry-based statistics, the requirement of analysing linked polymorphisms with occasional recombination means that values of *A_m_*are not always as straightforward to interpret (see below).

Like most asymmetry-based methods, our approach focuses on a very specific signal in WGS data to detect past gene flow. By contrast, model-based methods tend to use more comprehensive summaries of WGS data to infer past demography. Do model-based methods then implicitly use incongruent asymmetry to infer gene flow? Several popular demographic inference methods (Gutenkunst *et al*. 2009; Jouganous *et al*. 2017; Kamm *et al*. 2020; Excoffier *et al*. 2021) use the joint site-frequency spectrum (jSFS) as a convenient way to summarise polymorphism data from two or more populations. Because the jSFS only contains information about average branch lengths, it does not preserve direct information about incongruent asymmetry. Instead, the signal of gene flow in the jSFS is mostly contained in the sharing of low frequency derived alleles between populations (Gutenkunst *et al*. 2009), therefore requiring a large sample size. In contrast, *A_m_* requires only two diploid samples.

Other demographic inference methods leverage the joint distribution of branch lengths (Gronau *et al*. 2011; Flouri *et al*. 2023; Laetsch *et al*. 2023) and we therefore do expect these methods to implicitly use *A_m_* to infer rates of gene flow. Interestingly, we found that the distribution of *A_m_*values for the pair of *Heliconius* species did not match those expected under the demography inferred by Laetsch *et al*. (2023) (Figure 6A). This could be because their inference approach (which also uses the GF calculation from Lohse *et al*. 2011, 2016) assumes no recombination within a 64 bp sequence block (note that the expected value of *A_m_*for a block size of 1 bp under their demography is similar to observed *A_m_* for a 64 bp block in the data; Figure 6A). Alternatively, the difference could be because their method considers more than just incongruent asymmetry to infer gene flow. For example, the density of *fixed* polymorphisms provides information for inference under the approach of Laetsch *et al*. (2023), but is absent from *A_m_*. While the density of sequence polymorphisms does reflect past demography, it can also be influenced by variation in mutation rate across the genome. This has led to the development of three-taxon model-based methods that allow *µ* to vary between loci (Yang 2002, 2010; Galtier 2024). By contrast, *A_m_*-which relies on sampling from only two-taxa - is expected to be robust to such variation.

### Limitations

One strength of our approach is that we have an explicit expectation for *A_m_*under a history of strict isolation (*A_m_* = 0), whereas many related summary statistics instead require coalescent simulations to obtain expected values (Geneva *et al*. 2015; Rosenzweig *et al*. 2016; Hibbins and Hahn 2022). However, our analytic results only hold in the absence of recombination (Figure 4). We have circumvented this issue to some extent by calculating *A_m_* across varying block sizes, as some distributions provide unambiguous evidence for gene flow (Figures 4C and 6A). Our reanalysis of WGS data from *Odocoileus* deer, however, shows that certain demographic histories make detection of gene flow with *A_m_* challenging (Figure 6B). More specifically, when the difference in *N_e_* between populations is large, congruent genealogies tend to have highly asymmetrical external branch lengths (Figure 1C). As a result, blocks that span congruent genealogies due to hidden recombination events will have non-zero values of *A_m_*, even in the absence of migration. In the case of the *Odocoileus* deer it was helpful to perform coalescent simulations of previously inferred demographic histories (Kessler and Shafer 2024), as this suggested that *A_m_* is likely non-zero due to past gene flow, rather than due to differences in *N_e_* alone. Nevertheless, the influence of recombination on estimates of *A_m_* is certainly a weakness of our approach.

There are, in principle, several ways to remove the influence of recombination from estimates of *A_m_*. One option is to calculate *A_m_* (or *A_i_*) from the marginal genealogies of a reconstructed ARG (Rasmussen *et al*. 2014; Kelleher *et al*. 2019; Speidel *et al*. 2019). A perfectly inferred ARG would provide unbiased estimates of *A_m_*, yet it is unclear how often reconstructed marginal genealogies contain hidden recombination events, and whether integrating over a posterior sample of ARGs helps to resolve this issue. Either way, one could argue that estimating *A_m_* from an ARG is unnecessary, as it is already possible to infer much more detailed information about gene flow from such data (Wang *et al*. 2020; Pope *et al*. 2023). An alternative strategy is to derive expectations for *A_m_* in the presence of low rates of recombination, for example by including recombination in the GF (Lohse *et al*. 2011). Note, however, that analytic expectations would depend on all five parameters of the IM model as well as the rate of recombination. This means that estimation of *A_m_* would become a (challenging) model-based inference problem. There does not seem to be any way to easily adjust *A_m_* to be robust to recombination, and so careful interpretation of *A_m_* distributions (Figures 4 and 6) may be the most straightforward way to deal with this issue.

### Outlook

In this work we have shown that gene flow between two recently diverged populations leads to an asymmetry in genealogical branch lengths. Much of our focus has been on defining and exploring the properties of an asymmetry-based two-taxon test for gene flow. We do not, however, suggest that our test take the place of current model-based inference methods. Instead, we anticipate that the test will complement existing approaches, as it focuses on a particular signal that model-based methods either fail to capture or conflate with other information. More generally, we hope that our results provide some useful intuition about how gene flow shapes genealogical histories and genome sequence variation.

## Methods

### Coalescent simulations and block-jackknife resampling

We performed coalescent simulations to confirm our analytic results and to generate expectations of incongruent asymmetry under histories including recombination and bidirectional gene flow. We used msprime 1.3.0 to simulate histories under the IM model and tskit 0.5.6 to record the external branch lengths of genealogies (Baumdicker *et al*. 2021). In some cases, a simulated sequence corresponded to a single block, whereas in other cases a sequence contained multiple blocks. For each block we recorded values of *τ_a_* and *τ_b_* (or equivalently 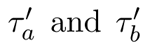). We summed *τ_a_* and *τ_b_* values across all blocks before calculating the scaled ratios *A_i_*and *A_m_*. We used a block-jackknife resampling procedure to estimate 95% confidence intervals (Quenouille 1949; Malinsky *et al*. 2021). To limit the computation time of the block-jackknife we grouped sequences blocks into jackknife-blocks (minimum of 25), ensuring that non-independent sequence blocks from the same simulation replicate were grouped together. Code for performing the coalescent simulations is provided as a python notebook (see Data availability).

### Analysis of real data

We reanalysed WGS data from *Heliconius* butterflies and *Odocoileus* deer. More specifically, we used the filtered VCF file from Laetsch *et al*. (2023) (shared via personal communication), and converted the genotypes from Kessler *et al*. (2023) (shared via personal communication) to a VCF file with the script genoToVCF.py (Martin and Amos 2021). We estimated *A_m_* across a range of block sizes from these VCF files using the script estimate Am.py (see Data availability). For the *Odocoileus* deer we analysed the individuals Ov ON6 and Oh WAI, as these are individuals with high sequencing coverage that were also used in the MSMC-IM analysis in Kessler *et al*. (2023). For the *Heliconius* butterflies we selected individuals ros.CJ2071.m and chi.CJ565.m as they were both male and had high sequence coverage. Jackknife blocks corresponded to 10^6^ consecutive SNPs for the *Odocoileus* deer (with blocks sometimes spanning multiple sequence scaffolds), and chromosomes for the *Heliconius* butterflies.

### Parametric bootstraps

We performed a parametric bootstrap analysis to gain further information about the results obtained in *Heliconious* butterflies and *Odocoileus* deer. Each replicate included simulation of a 66 kb sequence under a specific demography. The sequence was then split up into blocks ranging from 2^0^ (1) to 2^15^ (65, 536) bases in length. We used the same strategy as above to calculate point estimates and 95% confidence intervals of *A_m_*for each block size.

For the butterflies we performed 1000 replicate simulations under the IM demography of Laetsch *et al*. (2023). Here, *H. melpomene* (*N_mel_* = 549,000) and *H. cydno* (*N_cyd_* = 1,415,000) split from a common ancestor (*N_anc_* = 927,900) 4,216,000 generations ago. Backward in time, migration occurs from *H. mel* to *H. cyd* at rate *m_e_* = 7.4 *×* 10*^−^*^7^. Recombination occurs at rate *r* = 1.89 *×* 10*^−^*^8^ (Davey *et al*. 2017); mutation, at rate *µ* = 2.9 *×* 10*^−^*^9^ (Keightley *et al*. 2015) (see python notebook 2).

For the deer, we ran 1000 replicate simulations of the three-population demography from Figure 4a in Kessler and Shafer (2024), as well as 1000 replicates of the alternative demography in Figure 4b. Both models share the recombination rate *r* = 1.04 *×* 10*^−^*^8^ (Johnston *et al*. 2017) and mutation rate *µ* = 1.23 *×* 10*^−^*^8^ (Chen *et al*. 2019), as well as the property of bidirectional gene flow. The full parameters of these two demographic models are given in Table S5 of Kessler and Shafer (2024). Given their complexity, we do not describe the models here but code for simulating them can be found in python notebooks 3 and 4 (see Data availability).

## Data availability

Mathematica and python notebooks can be accessed at https://github.com/A-J-F-Mackintosh/ two_taxon_asymmetry. The Mathematica notebook is also available as a supplementary pdf file. The python script estimate Am.py can accessed at the same github repository (GNU General Public License v3.0). Sequence data from Laetsch *et al*. (2023) is available at the NCBI Sequence Read Archive (Bioprojects PRJEB11772 and PRJEB1749), as is the data associated with Kessler *et al*. (2023) (BioProject PRJNA830519).

## Acknowledgments

We would like to thank Meng Lu and Nicolas Galtier for providing feedback on a previous draft of this manuscript, as well as Konrad Lohse and Simon H. Martin for useful discussions. We would also like to thank Camille Kessler and Aaron B. A. Shafer for sharing the variant calls from Kessler *et al*. (2023), Dominik R. Laetsch for sharing the variant calls from Laetsch *et al*. (2023) and Bruna Cama for providing the *Heliconius* illustration used in Figure 6A. The *Odocoileus* illustration in Figure 6B is by Tracy A. Heath and was retrieved from PhyloPic (CC0).

## Funding

AM is supported by a grant from the Swedish Research Council (2022-03099) and DS is supported by a European Research Council (ERC) starting grant (ModelGenomLand 757648).

